# Comprehensive mutational scanning of a kinase *in vivo* reveals context-dependent fitness landscapes

**DOI:** 10.1101/004317

**Authors:** Alexandre Melnikov, Peter Rogov, Li Wang, Andreas Gnirke, Tarjei S. Mikkelsen

**Author notes:** Correspondence should be addressed to T. S. M.

## Abstract

Deep mutational scanning has emerged as a promising tool for mapping sequence-activity relationships in proteins^1–4^, RNA^5^ and DNA^6–8^. In this approach, diverse variants of a sequence of interest are first ranked according to their activities in a relevant pooled assay, and this ranking is then used to infer the shape of the fitness landscape around the wild-type sequence. Little is currently know, however, about the degree to which such fitness landscapes are dependent on the specific assay conditions from which they are inferred. To explore this issue, we performed deep mutational scanning of APH(3’)II, a Tn5 transposon-derived kinase that confers resistance to aminoglycoside antibiotics^9^, in *E. coli* under selection with each of six structurally diverse antibiotics at a range of inhibitory concentrations. We found that the resulting fitness landscapes showed significant dependence on both antibiotic structure and concentration. This shows that the notion of essential amino acid residues is context-dependent, but also that this dependence can be exploited to guide protein engineering. Specifically, we found that differential analysis of fitness landscapes allowed us to generate synthetic APH(3’)II variants with orthogonal substrate specificities.

---

To enable efficient generation of comprehensive and relatively unbiased, single amino acid substitution libraries for deep mutational scanning and related applications, we developed a highly multiplexed approach to site-directed mutagenesis that we refer to as Mutagenesis by Integrated TilEs (MITE; Fig. 1). Briefly, we design a set of long DNA oligonucleotides that encode all desired mutations, but are otherwise homologous to the template sequence. The oligonucleotides are organized into ‘tiles’, where those within each tile differ in a central variable region but share identical 5’ and 3’ ends. The tiles are staggered such that their variable regions collectively span the entire template. The oligonucleotides are synthesized on a programmable microarray^10^ and individual tiles are then PCR amplified using primers complementary to their shared ends. To avoid hybridization and extension of partially overlapping oligonucleotides, the tiles can be split into two non-overlapping pools that are synthesized and amplified separately. Finally, the PCR products from each tile are inserted into linearized plasmids that carry the remaining template sequence using multiplexed sequence- and ligation-independent cloning. Because only a portion of each resulting mutant sequence is derived from the oligonucleotide pool, this approach limits the impact of synthesis errors but still benefits from the cost efficiency of microarray-based DNA synthesis.

**Figure 1.**
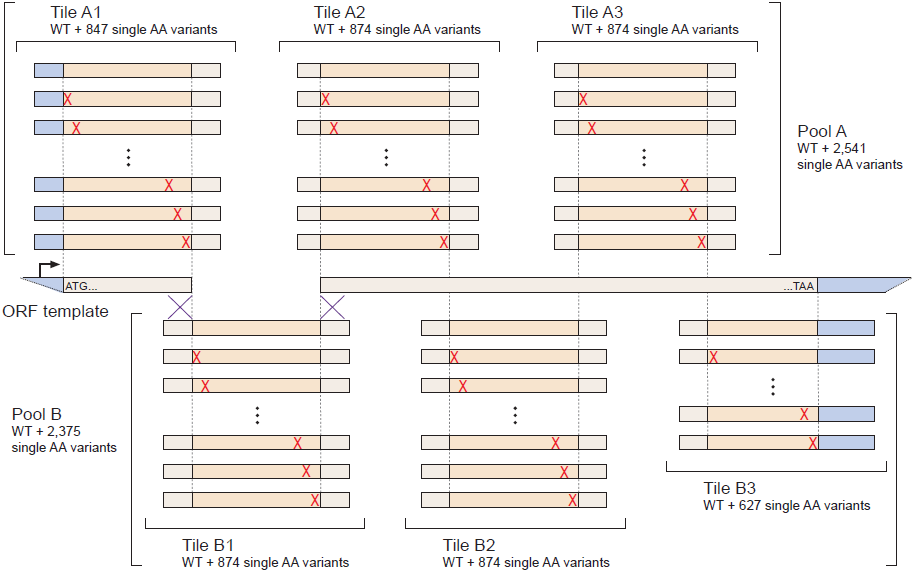
Overview of MITE. Oligonucleotides encoding all desired amino acid substitutions in a template open reading frame (ORF) are synthesized in two pools. The oligonucleotides in each pool are organized into non-overlapping tiles with shared 5’- and 3’ ends that facilitate selective PCR amplification. After amplification, each tile is inserted into a linearized expression vector that contains the remainder of the ORF using sequence- and ligation-independent cloning techniques.

We next applied MITE to perform deep mutational scanning of the Tn5 transposon-derived aminoglycoside-3’-phosphotransferase-II (APH(3’)II), a 264 amino acid residues kinase that confers resistance to a variety of aminoglycoside antibiotics^9^, with the goal of elucidating which residues are essential for its activity and whether these residues are the same for different substrates. We first designed and synthesized six 200 nucleotide (nt) tiles that encoded all possible single amino acid substitutions across APH(3’)II. Each tile contained a 140 nt variable region flanked by 30 nt constant ends. To test whether the cost-efficiency of our method could be further improved by synthesizing multiple mutant libraries in parallel, we also included corresponding tiles for nine homologous proteins, resulting in two non-overlapping pools of 26,250 and 23,666 distinct oligonucleotide sequences (Supplementary Data 1). We found that all six APH(3’)II tiles could be selectively amplified from these pools with minimal optimization of PCR conditions. We then inserted these tiles into plasmids that carried the corresponding remainders of the APH(3’)II coding sequence. Each amplification and multiplexed cloning reaction was performed in duplicate to generate two mutant libraries.

To characterize the resulting libraries, we first shotgun-sequenced their APH(3’)II coding regions to a depth of ∼120,000X (Supplementary Fig. 1). At this depth, we could detect 4,993 of 4,997 possible substitutions and found that the frequency of each mutant amino acid at each position was close to the expected value from an ideal single substitution library (observed median = 0.016%, interquartile range = 0.011%-0.023%, expected = 0.020%). Amino acid mutations caused by single nucleotide substitutions were slightly overrepresented, which is likely due to the combined effects of synthesis- and sequencing-related errors. We next sub-cloned 90 plasmids from the libraries for full-length sequencing of the coding region and found that the majority (90%) of these clones carried a single amino acid substitution, while the rest carried either none (2.2%) or two substitutions (7.8%). To ensure full representation and minimize the impact of synthesis errors, each library was titrated to contain plasmids from ∼10^7^ transformants.

To perform mutational scanning *in vivo*, we cultured *E. coli* transformed with APH(3’)II substitution libraries in liquid media supplemented with decreasing concentrations of one of six aminoglycoside antibiotics with diverse structures and a wide range of potencies: kanamycin, ribostamycin, G418, amikacin, neomycin or paromomycin (Supplementary Fig. 2). To normalize the selective pressures induced by the different antibiotics, we first established their minimum inhibitory concentrations (MICs) for *E. coli* transformed with “wild-type” (WT) APH(3’)II and then used 1:1, 1:2, 1:4 and 1:8 dilutions of WT MICs for the respective library selections. We note that, with the exception of G418 and amikacin, even the 1:8 dilutions were sufficient to inhibit growth of untransformed *E. coli*. After overnight selection, plasmids were isolated from the surviving cells and their coding regions were shotgun sequenced to compare the amino acid composition of the selected libraries to that of the original input libraries (median coverage = 51,400X; Supplementary Data 2).

We began our analysis by examining changes in the relative abundance of mutant versus WT amino acids at each position after selection with kanamycin (Fig. 2a). At the highest concentration (1:1 WT MIC), we observed a significant depletion of mutant amino acids at approximately 210 of 263 positions (χ^2^-test at the 5% false discovery rate (FDR) threshold, as determined by the Benjamini-Hochberg procedure^12^). At lower concentrations, we observed both a decrease in the number of positions with significant depletion of mutations (approximately 155, 130 and 113 at 1:2, 1:4 and 1:8 WT MIC, respectively; 5% FDR) and an increase in the diversity of amino acids that were tolerated at the remaining positions. Conservative substitutions to amino acids with biochemical properties that were similar to the WT residue were on average ∼1.4-fold more likely to be tolerated compared to non-conservative substitutions^13^, regardless of the kanamycin concentration (Supplementary Fig. 3a). The tolerance for specific substitutions appeared, however, to be both concentration- and position-dependent. Importantly, the observed changes in amino acid frequencies were highly consistent between independent selections (Pearson’s r = 0.76-0.91 across the four kanamycin dilutions; Supplementary Fig. 4).

**Figure 2.**
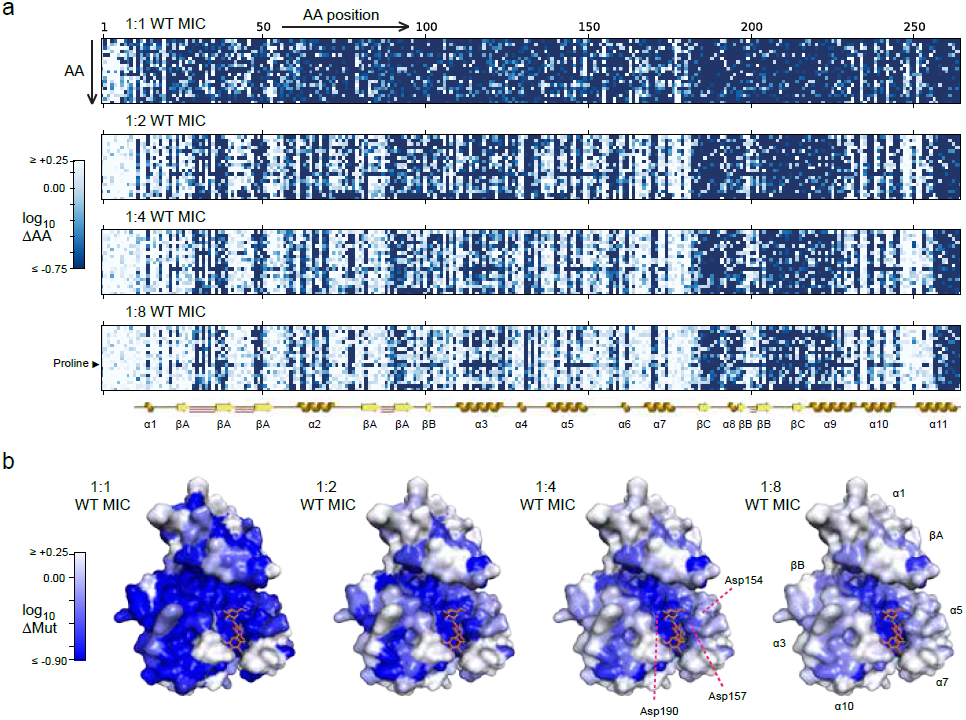
Mutational scanning of APH(3’)II using kanamycin selection. (**a**) Visual representation of the changes in abundance of each amino acid (ΔAA) at each position after selection at four different concentrations. The color in each matrix entry corresponds to the change relative to the input library. The known secondary structure of APH(3’)II is shown for reference, including alpha helices (gold), beta sheet strands (yellow) and beta hairpins (red; the first 10 residues are unstructured). The highly deleterious effect of proline substitutions, which impose unique structural constrains, stand out at low concentrations. (**b**) Projections of the observed changes in the abundance of mutant amino acids (ΔMut) onto a crystal structure of APH(‘3)II in complex with kanamycin (PDB accession: 1ND4). Numerical values are provided in Supplementary Data 2.

To better understand the patterns of selection, we projected them onto a previously reported co-crystal structure of APH(3’)II in complex with kanamycin^9^ (Fig. 2b). These projections show that while mutations are depleted from positions all over the protein at high concentrations, the positions that do not tolerate mutations even at low concentrations tend to cluster internally and near the active site of the enzyme. The projections also provide insights into selection patterns at specific positions. For example, residues 190, 157 and 154 are all aspartates in the vicinity of the active site in the WT enzyme (Supplementary Fig. 3b). At 1:1 WT MIC, we observed depletion of all amino acids except the wild-type aspartates at 190 and 157, while both aspartate and glycine were tolerated at 154. At more moderate selection pressures (1:4 WT MIC), aspartate remained the only amino acid tolerated at 190, aspartate and asparagine were both tolerated at 157, while a wide variety of amino acids, including glycine, valine and arginine, were tolerated at 154. Consistent with its apparent immutability, structural and functional studies have previously identified Asp190 as the catalytic residue in APH(3’)II and it is conserved across all known aminoglycoside phosphotransferases^9^. The crystal structure also shows that Asp157 forms a stabilizing hydrogen bond with the second ring of kanamycin, which explains why substitutions to amino acids other than the very similar asparginine might be deleterious. In contrast, Asp154 does not appear to directly interact with the substrate, which is consistent with the higher tolerance we observed for non-conservative substitutions at this position. These results support the quantitative and qualitative accuracy of our mutational scanning data, but also suggest that the distribution of relative fitness among the mutants is highly sensitive to the conditions in which they are assayed. We note that if this is a general phenomenon, then establishing quantitative control over selection pressures will be critical to interpretation of deep mutational scanning experiments.

To compare the selection patterns induced by kanamycin in our experiments to those that have molded APH(3’)II over evolutionary timescales, we examined a conservation profile derived from alignment of 133 homologs^14^. We found positive rank correlations between evolutionary conservation and depletion of mutant amino acids at most positions across the protein (from Spearman’s ρ = 0.51 at ∼1:1 WT MIC to ρ = 0.74 at 1:8 WT MIC; Supplementary Fig. 5a). The most common discrepancies were positions that were relatively conserved, yet highly tolerant of mutations under selection with kanamycin. For example, we identified 11 conserved residues that consistently tolerated non-WT amino acids even at 1:1 WT MIC (Supplementary Fig. 5b). Interestingly, 7 of the 11 residues directly flank the active site, but none of them appear to interact with kanamycin in the WT enzyme. This may reflect that kanamycin is not the substrate, or at least not the only substrate, that has driven the natural evolution of APH(3’)II, and consequently that artificial selection with kanamycin will tolerate or even favor some substitutions that would be deleterious in other contexts.

We next examined the effects of selection using the five other aminoglycosides. These substrates generated concentration- and position-dependent selection patterns that were qualitatively similar to those generated by kanamycin (Supplementary Figs. 6-10). To quantitatively compare the patterns, we ranked the twenty amino acids by the relative changes in their abundances after selection and then computed the rank correlations at each position for each pair of conditions. We found that selection with any two aminoglycosides at matched concentrations showed positive correlations at the majority of positions (for example, median Spearman’s ρ = 0.79 for kanamycin and paromomycin at 1:2 WT MIC and ρ > 0 at 235 of 264 positions at 5% FDR; Supplementary Fig. 11), which implies that the relative fitness values of the mutants were largely similar. We did, however, notice some substitutions that were consistently tolerated under selection with one substrate but depleted under selection with another. We hypothesized that further analysis of these exceptions might identify residues that influence the relative activity of APH(3’)II on different substrates.

To identify such specificity-determining residues, we queried our data for individual substitutions that showed significant depletion after kanamycin selection at 1:2 WT MIC (at 5% FDR) but no trend towards depletion after selection with a second aminoglycoside at its highest concentration. These stringent criteria identified a handful of substitutions that appeared to be well-tolerated in the presence of ribostamycin, G418 or paromomycin but not kanamycin (11, 3 and 4 single substitutions, respectively; Supplementary Fig. 12a; Supplementary Table 1). To confirm, we re-synthesized APH(3’)II variants containing a selection of these substitutions and compared their specificities towards the second aminoglycoside to that of WT APH(3’)II (here, we define specificity as the ratio of the MIC of the second aminoglycoside over the MIC of kanamycin). In six out of nine cases, the tested substitution led to a ≥2-fold increase in specificity towards the second aminoglycoside (range: 2- to 8-fold; see Supplementary Table 2). Moreover, in nine of eleven additional test cases, we found that combining two or more substitutions that favored the same aminoglycoside increased the specificity to a greater extent than any one of the substitutions did alone (range: 4- to 32-fold; Supplementary Table 2). This indicates that our approach accurately identified specificity-determining residues in APH(3’)II and that these residues can have additive effects.

To expand our search for specificity-determining residues, we next looked for substitutions that showed significant depletion after kanamycin selection at 1:2 WT MIC (at 5% FDR) but no trend towards depletion by the second aminoglycoside at its 1:2 WT MIC, or *vice versa*. These less stringent criteria identified additional candidates in every pairwise comparison (Fig. 3a-e; Supplementary Table 1). Many of these candidate substitutions flanked the active site, particularly at or near residues that interact with the second and third ring of kanamycin in the WT crystal structure (in particular, the α5-α7 and α9-α10 helices). This has been reported as an expected hotspot for specificity-determining residues, because the first ring is constrained by being the phosphate acceptor, while the remaining rings show significant structural diversity among the aminoglycosides^9^. Candidate substitutions were also frequently found in a relatively flexible part of the N-terminal domain that extends over the phosphate donor (ATP) and aminoglycoside binding sites (in particular, positions 27-57), and in the linker region between the N-terminal domain and the central core (in particular, position 96). These substitutions may influence the presentation of the two substrates near the catalytic residue, but the absence of ATP in the crystal structure limits potential insights from the available model. To confirm the expected functional effects of the identified substitutions, we synthesized and determined the substrate-specificity of five pairs of designed APH(3’)II variants, where each pair incorporated all substitutions that were preferentially tolerated by either kanamycin or the second aminoglycoside in the respective comparison (if multiple candidate substitutions were found at one position, we picked the one that showed the largest difference in abundance). Nine of the ten tested variants showed changes in specificity that favored the targeted substrate (range: 2- to 128-fold; Fig. 3f and Supplementary Table 2). The tenth variant had minimal activity on both substrates, which was likely a consequence of the relatively large number of substitutions in this variant (24 compared to 12 or less in the others). Indeed, in six of the nine successful cases, the expected increase in specificity also came with a concomitant but lesser decrease in absolute activity (as measured by the relevant MIC; Supplementary Table 2), which suggests a precarious balance between the two properties. Strikingly, the two variants that were designed to favor or disfavor paromomycin relative to kanamycin showed a >2,000-fold combined change in their specificities. In this case, a single round of MITE-enabled mutational scanning was therefore sufficient to derive two synthetic enzyme variants with essentially orthogonal substrate specificities from one non-specific progenitor (Fig. 3g).

**Figure 3.**
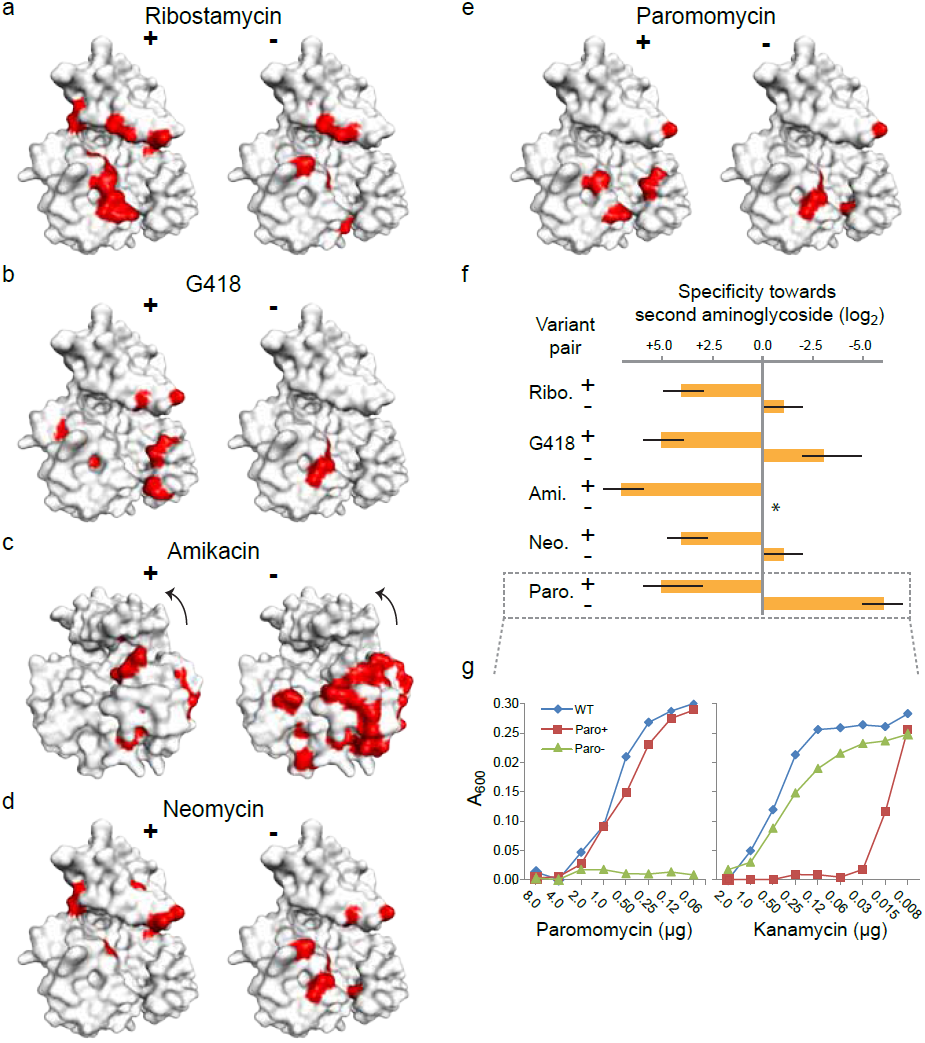
Identification of specificity-determining residues in APH(3’)II. (**a**)-(**e**) The positions of substitutions that are specifically tolerated (+) or depleted (-) under selection with the indicated aminoglycoside relative to selection with kanamycin are shown in red. The identities of the substitutions are listed in Supplementary Tables 1 and 2. Arrows indicate a rotation of the structure. (**f**) Specificity towards the indicated aminoglycoside for pairs of synthetic enzyme variants designed to favor (+) or disfavor (-) this substrate over kanamycin, relative to that of WT APH(‘3)II. Bars show the medians and error bars show the ranges observed over 2-3 independent cultures, bounded by the resolution of the assay (2-fold dilutions; see also Supplementary Table 2). The variant predicted to favor kanamycin over amikacin (Ami. -) showed minimal activity on both substrates. (**g**) Optical density in *E. coli* cultures transformed with WT APH(3’) or synthetic variants designed to specifically favor paromomycin (Paro+) or kanamycin (Paro-), after selection with each of these two aminoglycosides.

In summary, MITE coupled with deep sequencing allowed us to perform quantitative mutational scanning of the complete APH(3’)II protein coding sequence with sufficient accuracy to identify individual kinase activity- and substrate specificity-determining residues. We emphasize that structural information or molecular dynamics simulations was not required or directly utilized to identify these residues. Our approach is therefore applicable even when accurate structural models are unavailable or intractable. Assuming that indirect selection strategies can be devised, it can also be extended to proteins and activities that do not confer a direct growth advantage. Notably, we have recently used a similar approach to identify activity- and specificity-determining nucleotides in mammalian gene regulatory elements^8^. In both cases, we found that combining mutational scanning data from two separate conditions is an effective method for identifying mutations that change the ratio of the activities in the two conditions in the desired direction. Engineering families of biomolecules with orthogonal activities is a central challenge in biotechnology and synthetic biology^2,15^, but traditional directed evolution is poorly suited to this task due to the difficulty of performing direct selection for the absence of activity in one or more conditions. We therefore expect that performing parallel assays of designed mutant libraries under multiple conditions, followed by integrative analysis and iterative synthesis of new variants, will be a useful strategy for dissecting and engineering a wide variety of biomolecular activities and interactions.

## METHODS

### Plasmid construction and cloning

All sequences used in this study, including gene synthesis products, oligonucleotide libraries and primers, are provided as Supplementary Data 1. Primers and individual oligonucleotides were synthesized by Integrated DNA Technologies (Coralville, IA), the two 200mer oligonucleotide pools containing mutagenesis tiles were synthesized by Agilent (Santa Clara, CA) as previously described^10^, the WT APH(3’)II coding region template for library construction was synthesized by GenScript USA (Piscataway, NJ), and the single- and multi-substitution synthetic ORF variants for validation of specificity-determining amino acid residues were synthesized by Gen9 (Cambridge, MA).

To generate a plasmid vector carrying a constitutive EM7 promoter (Life Technologies) and a synthetic gene encoding APH(‘3)II (*neo*; UniProtKB entry name: KKA2_KLEPN) flanked by two *SacI* restriction sites, we first combined a DNA fragment containing EM7 (template: EM7_pBR322; primers: EM7_Amp_F/R) with a PCR-linearized pBR322 backbone (template: pBR322; primers: pBR322_EM7_F/R) using the In-Fusion PCR Cloning System (Clontech Laboratories, Mountain View, CA). The resulting plasmid (pBR322[EM7]) was then re-linearized by inverse PCR (primers: pBR322_tetRLin_F/R) and combined with a PCR amplified *neo* ORF fragment (template: KKA2_KLEPN_opt; primers: KKA2_pBR322_F/R) by In-Fusion to replace the *tetR* ORF of pBR322 with *neo*. The final product (pBR322[EM7-*neo*]) was isolated using QIAprep Spin Miniprep kits (Qiagen, Gaitehrsburg, MD) and verified by Sanger sequencing (primers: pBR322_Seq_F/R).

To generate the APH(‘3)II single-substitution libraries, full-length oligonucleotides were first isolated from the synthesized pools using 10% TBE-Urea polyacrylamide gels (Life Technologies, Carlsbad, CA). Each pair of APH(3’)II tiles and corresponding linearized pBR322[EM7-*neo*] plasmids were PCR amplified by Herculase II DNA Polymerase (Agilent) with their respective primers (primer prefix: KKA2_TileAmp for tiles, KKA2_LinAmp for backbones), size selected on 1% E-Gel EX agarose gels (Life Technologies), purified with MinElute Gel Extraction kits (Qiagen) and combined using In-Fusion reactions. Each reaction was separately transformed into Stellar chemically competent cells (Clontech), the transformants were grown in LB media with carbenicillin (50 µg/mL) and plasmid DNA libraries were isolated using QIAprep Spin Miniprep kits (Qiagen). Finally, complete substitution libraries were generated by pooling equimolar amounts of the resulting six single-tile plasmid libraries.

To generate plasmids encoding selected single- and multiple-substitution APH(3’)II variants, we amplified the Gen9 synthetic ORFs using Herculase II DNA Polymerase (primers: KA2_pBR322_F/R) and inserted them into a PCR-linearized backbone (primers: pBR322_tetLin_F/R, template: pBR322[EM7]) using In-Fusion reactions. The resulting plasmids (pBR322[EM7-K1] through pBR322[EM7-K44]) were verified by Sanger sequencing and preserved in *E. coli* as glycerol stock.

### Cell culture and selection

To determine the minimum inhibitory concentration (MIC) of each of the six aminoglycosides: kanamycin, ribostamycin, G418, amikacin, neomycin, paromomycin (Sigma-Aldrich, St. Louis, MO) in *E. coli* expressing WT APH(3’)II, Stellar cells were transformed with pBR322[EM7-*neo*], recovered in SOC medium (New England Biolabs (NEB), Ipswich, MA) at 37°C for 1 hr, diluted 1:100 with LB and then divided into 96-well growth blocks containing LB with carbenicillin (50 µg/mL) and 2-fold serial dilutions of aminoglycosides. After growth at 37°C with shaking for 24 hrs, the culture densities were assessed by absorbance at 600 nm (A_600_) using a NanoDrop 8000 (Thermo Scientific, Billerica, MA). The MIC for each aminoglycoside was estimated as the lowest dilution at which A600 was less than 0.025.

To perform mutational scanning, Stellar cells were transformed with 10 ng (∼3 fmol) of mutant library plasmids, recovered in SOC medium at 37°C for 1 hour, diluted into 15 mL LB with carbenicillin (50 µg/mL) and one of the aminoglycosides at 1:1, 1:2, 1:4 or 1:8 dilutions of the following concentrations (estimated 1:1 WT MICs): 225 µg/mL for kanamycin, 2500 µg/mL for ribostamycin, 5 µg/mL for G418, 10 µg/mL for amikacin (see notes in Supplementary Data 1), 40 µg/mL for neomycin and 320 µg/mL for paromomycin. The cultures were incubated in 50 mL tubes at 37°C with shaking for 24 hours, pelleted by centrifugation and frozen at −20°C. Plasmids were isolated from the pellets using QIAprep Spin Miniprep kits (Qiagen). Each transformation and selection was performed in duplicate, using each of the two independently generated mutant libraries.

To establish the substrate specificity of the synthetic APH(3’)II variants relative to WT APH(3’)II, glycerol stocks of the relevant clones (pBR322[EM7-K1] through pBR322[EM7-K44]) were streaked onto LB agar plates and cultured overnight at 37°C. Single colonies were then inoculated into 1 mL LB with 50 µg carbenicillin, incubated at 37°C with shaking for 6 hours, diluted 1:100 into LB media, and then split into 96-well deep well plates containing LB with carbenicillin (50 µg/mL) and 2-fold serial dilutions of aminoglycosides dilution series. We note that the longer recovery in these follow-up experiments increased the absolute MICs compared to those we measured immediately following transformation).

### Mut-Seq

To sequence and count mutations, the APH(3’)II coding regions were first isolated from the plasmid pools by *SacI* digest (NEB) followed by agarose gel purification. The *SacI* fragments were ligated into high molecular weight concatemers using T4 DNA ligase (NEB). The concatemers were fragmented and converted to sequencing libraries using Nextera DNA Sample Prep kits (Illumina, San Diego CA). Library fragments from the 200-800 nt size range were selected using agarose gels and then sequenced on Illumina MiSeq instruments using 2×150 nt reads. The reads were subsequently aligned to a reference sequence consisting of concatenated copies of the WT APH(3’)II coding region using BWA version 0.5.9-r16 with default parameters^16^. The number of occurrences of each amino acid at each position along the coding region was then counted and tabulated from the aligned reads. Only codons for which all three sequenced and aligned nucleotides had phred quality scores ≥ 30 were included in these counts.

### Computational analysis

Data processing and statistical analysis, including computation of Pearson’s and Spearman’s correlation coefficients, χ^2^-statistics and associated p-values, were performed using the Enthought Python Distribution (http://www.enthought.com), with IPython^17^, NumPy version 1.6.1, SciPy version 0.10.1 and matplotlib version 1.1.0^18^. Rendering of the APH(3’)II crystal structure (PDB accession 1ND4) was performed using PyMOL version 1.5 (Schrödinger). The rendering of its secondary structure was derived from PDBsum at EMBL-EBI (http://www.ebi.ac.uk/pdbsum/). The evolutionary conservation profile for APH(3’)II was obtained from ConSurf-DB^14^.

The magnitude of changes in the abundance of mutant amino acids at each position after selection was estimated as 

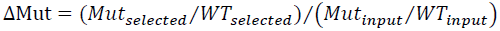

where *Mut*_*input*_ and *WT*_*input*_ are, respectively, the observed counts of mutant and wild-type amino acids at that position in the input library and *Mut*_*selected*_ and *WT*_*selected*_ are the corresponding observed counts after selection.

The magnitude of changes in the abundance of each specific amino acid at each position after selection was estimated as 

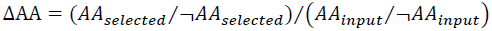

where *AA*_*input*_ and ¬*AA*_*input*_ are, respectively, the observed counts of that amino acid and all other amino acids at that position in the input library and *AA*_*selected*_ and ¬*AA*_*selected*_ are the corresponding observed counts after selection.

The statistical significance of a deviation of any ΔMut or ΔAA from 1.0 was estimated using a χ^2^-test for independence on a 2×2 contingency table that contained the four corresponding counts with a pseudocount of 1 added to each. To correct for multiple hypothesis testing, the Benjamini-Hochberg procedure was applied to identify the 5% false discovery rate (FDR) threshold^12^. We note that ΔMut or ΔAA values greater than 1.0 do not necessarily indicate higher fitness relative to the WT APH(3’)II (i.e., positive selection), because the majority of ORFs that contributed to the count of WT residues at one position still carried substitutions at other positions.

Substitutions that favored growth under selection with one aminoglycoside relative to another were identified by requiring ΔAA < 1.0 at 5% FDR across two replicates under selection with the first and ΔAA ≥ 1.0 across two replicates for the same substitution under selection with the other, as well as a minimum difference of 0.5 between the log_10_-transformed ΔAA values, at the concentrations indicated in the main text. These thresholds were established empirically to select a limited number of high confidence candidates.

## ACKNOWLEDGEMENTS

The authors would like to thank E. M. LeProust and S. Chen for oligonucleotide library synthesis, and X. Zhang, A. Majithia, G. S. Ramachandran, members of the Broad Technology Labs and the members of the Lander laboratory for discussions. This project was supported by funds from the Broad Institute.

## AUTHOR CONTRIBUTIONS

A.M. and T.S.M. developed the MITE method and designed the study. A. M., P. R., A. G. and T. S. M. performed or supervised the molecular biology experiments. L. W. and T. S. M. performed the cell culture experiments. T.S.M. analyzed the data and wrote the manuscript with substantial input from all authors.

## ACCESSION NUMBERS

All sequencing data has been submitted to the NCBI Short Read Archive under accession [pending].

---

**Supplementary Figure 1.**
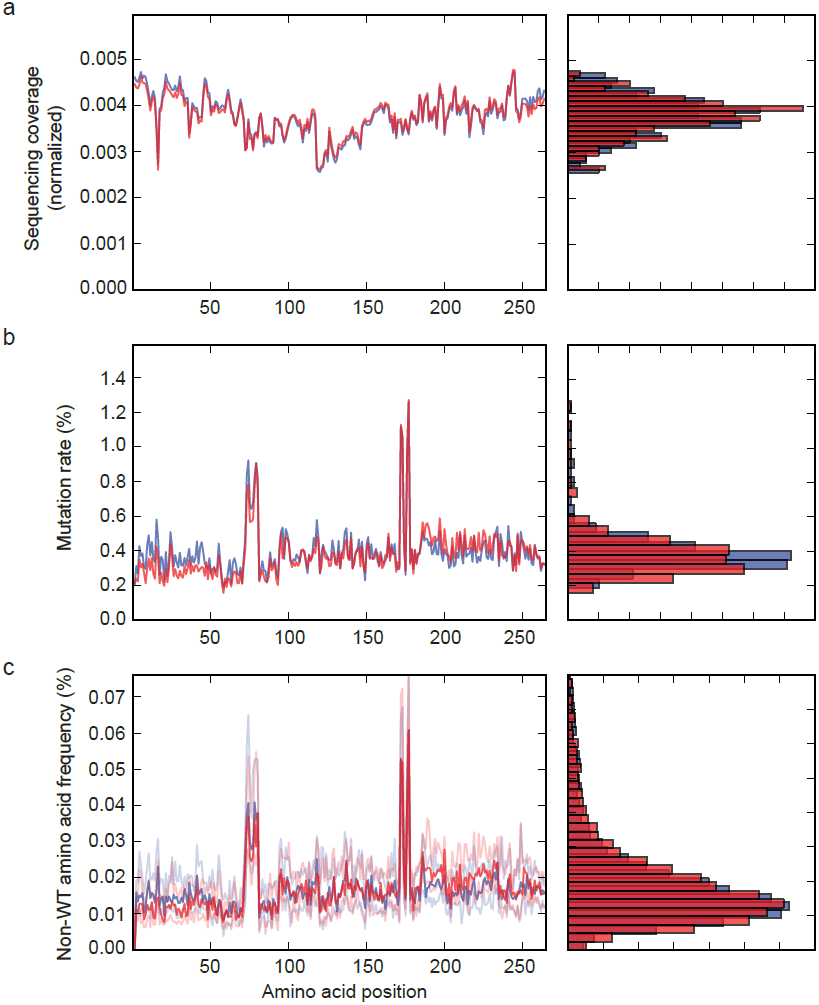
Sequencing-based quality control of the APH(’3)II mutant libraries prior to selection. Two libraries (red, blue) were independently constructed from the same raw oligonucleotide synthesis products. a. High-quality sequence coverage across the APH(3’)II coding region, as represented by the number of observations of each codon with phred quality scores ≥ 30 at all three positions (normalized by division by the median raw count). The observed variation is likely caused by insertional bias of the Nextera transposons.
b. Mutation rate (# of non-WT amino acids/# of amino acids observed per position). Two hotspots appear to trace back to overrepresentation of a subset of oligonucleotides in the raw synthesis product, but their effects on the overall library composition are limited.
c. Frequencies of each non-WT amino acid (# of one amino acid/# of all amino acids) at each position. The dark lines in the left-hand side plot show the median and the light lines show the 1st and 3rd quartiles at each position. The right-hand side histograms show the complete distribution (n=4,997).

**Supplementary Figure 2.**
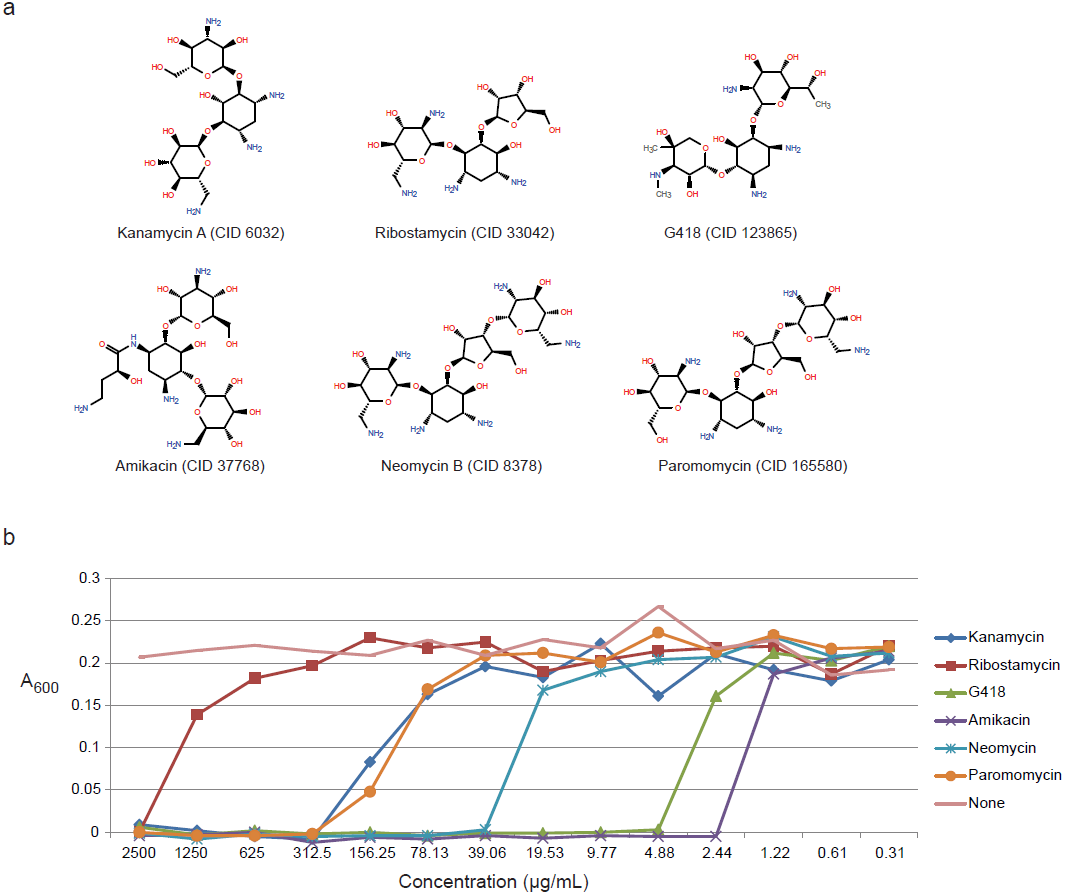
a. 2-dimensional structures of the six aminoglycoside antibiotics used for selection in this study. The structures were obtained from PubChem (http://pubchem.ncbi.nlm.nih.gov/) and rendered using Open Babel (http://openbabel.org/).
b. Optical density (600 nm) in cultures of *E. coli* transformed with WT pBR322[EM7-*neo*] after 24 hours in liquid LB supplemented with the indicated antibiotic concentrations and 50 μg/mL carbencillin. The potency of the six compounds, as estimated by their minimum inhibitory concentrations (MICs), differ by three orders of magnitude.

**Supplementary Figure 3.**
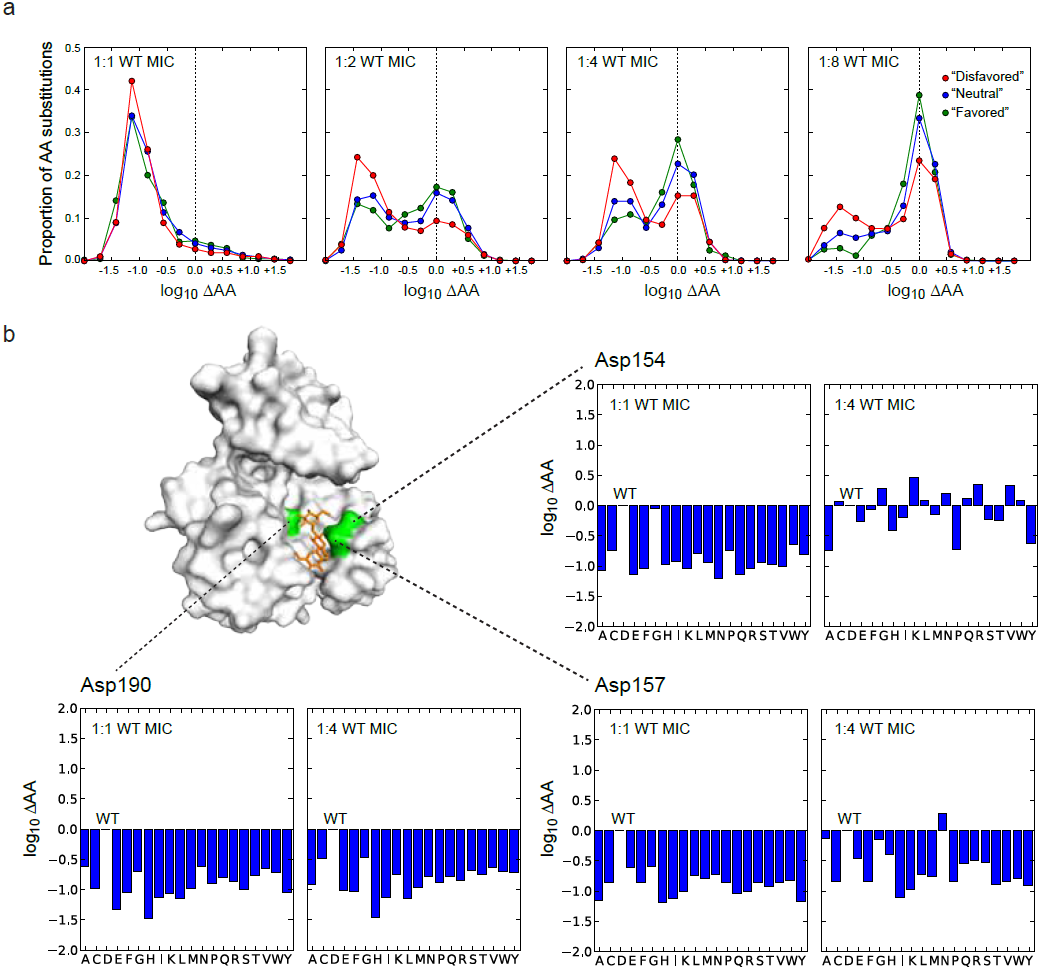
a. Distributions of changes in amino acid frequencies (ΔAA) at each position after selection with kanamycin at four concentrations. The amino acids are divided into three classes, according to whether they would be considered a “disfavored”, “neutral” or “favored” substitution from the WT residue in an intracellular protein purely based on their biochemical properties (as estimated by Russell et al, http://http://www.russelllab.org/aas/, accessed April 2013). A greater diversity of substitutions were tolerated at low concentrations, although the ratio of “favored” over “disfavored” substitutions that were well-tolerated (log_10_ ΔAA ≥ 0.0) was relatively constant at ∼1.4.
b. Examples of concentration- and position-dependent selection patterns for three positions containing aspartate resiudes in the WT APH(3’)II protein. The bar plots show the changes in the observed frequency of each amino acid after selection at the indicated concentrations. Asp190 is believed to be the primary catalytic residue in APH(3’)II, while Asp157 contributes a hydrogen bond in the APH(3’)II-kanamycin co-crystal.

**Supplementary Figure 4.**
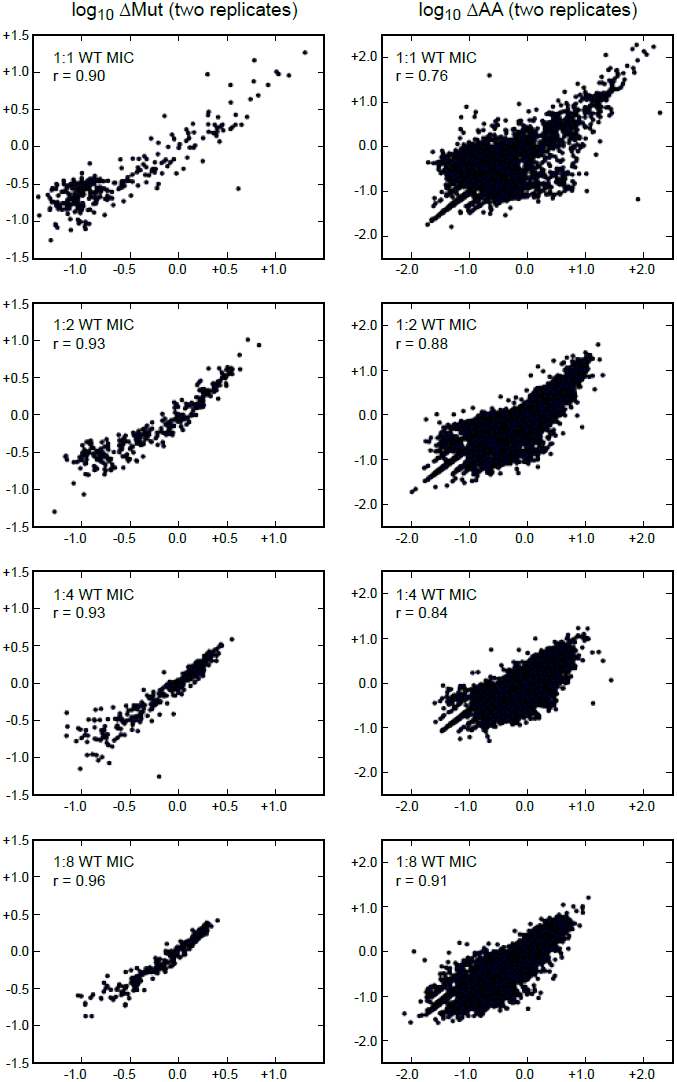
Scatter plots of the observed changes in the frequencies of mutant amino acids (left) or individual amino acids (right) in two independent kanamycin selection experiments at the indicated concentrations. The Pearson correlation coefficients (r) are all significant at p < 10^-20^.

**Supplementary Figure 5.**
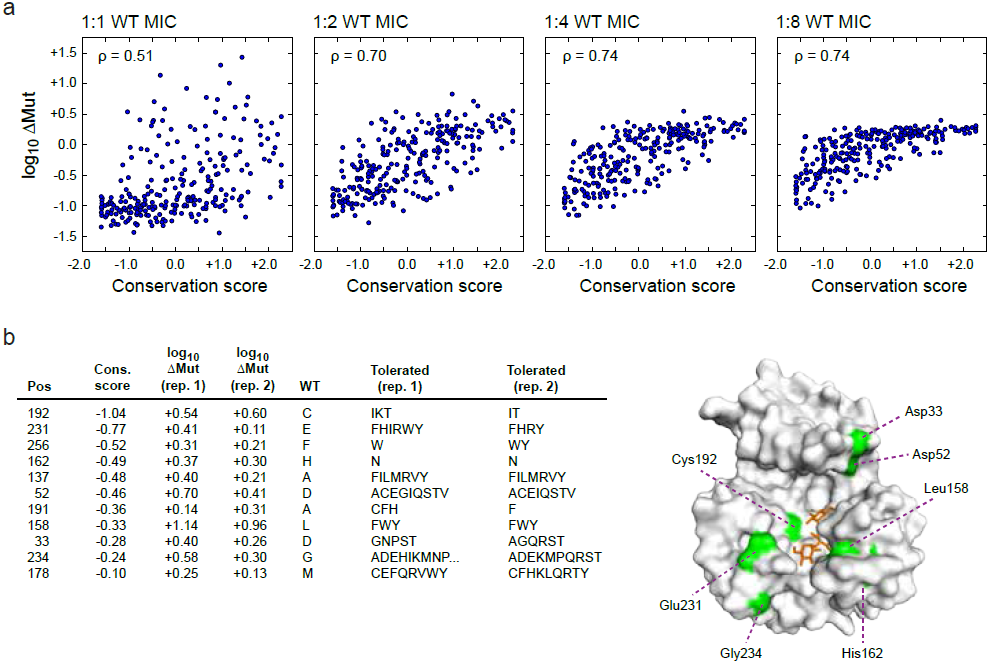
a. Scatter plots and Spearman’s rank correlation coefficients comparing the level of evolutionary conservation at each position along APH(3’)II to the observed changes in the frequencies of mutant amino acids under selection with kanamycin at the indicated concentrations. The conservation scores were obtained for the from ConSurf-DB (http://bental.tau.ac.il/new_ConSurfDB/, accession 1ND4, April 2013).
b. Enumeration of the positions that show relatively strong evolutionary conservation (conservation score < 0.0), but also high tolerances for one or more non-WT substitutions (log_10_ ΔMut > 0.0) under selection with kanamycin at 1:1 WT MIC across two replicates.

**Supplementary Figure 6.**
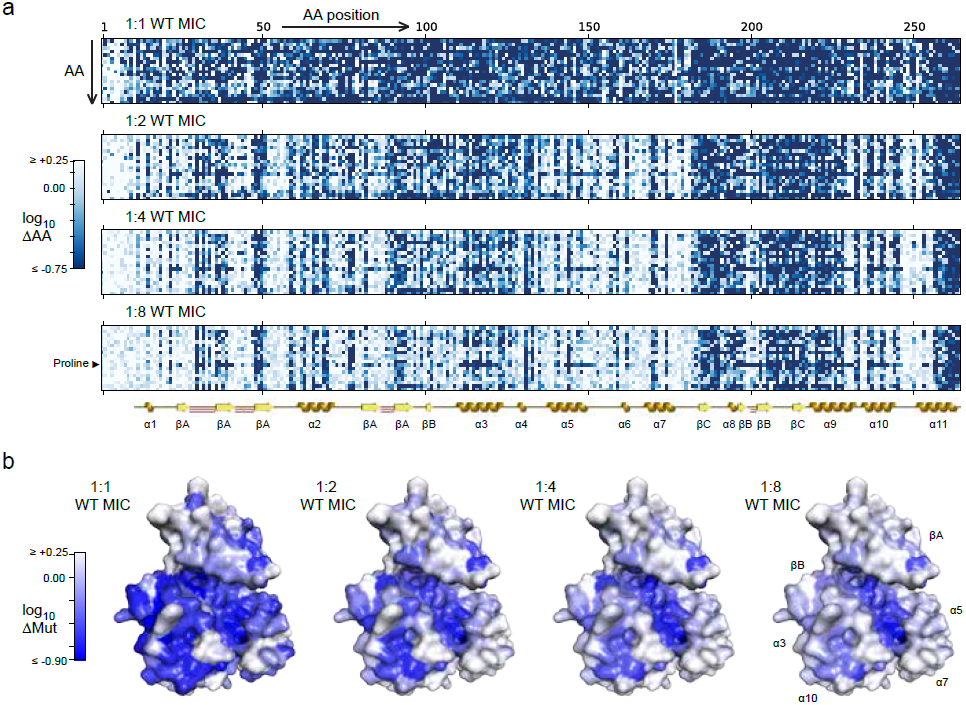
Mutational scanning of APH(3’)II using ribostamycin selection. a. Visual representation of the abundance of each amino acid (ΔAA) at each position after selection at four different concentrations. The color in each matrix entry corresponds to the change relative to the input library. The known secondary structure of APH(3’)II is shown for reference, including alpha helices (gold), beta sheet strands (yellow) and beta hairpins (red; the first 10 residues are unstructured).
b. Projections of the observed changes in the abundance of mutant amino acids (ΔMut) onto a crystal structure of APH(‘3)II in complex with kanamycin (PDB accession: 1ND4; the kanamycin substrate is not show). Numerical values are provided in Supplementary Data 2.

**Supplementary Figure 7.**
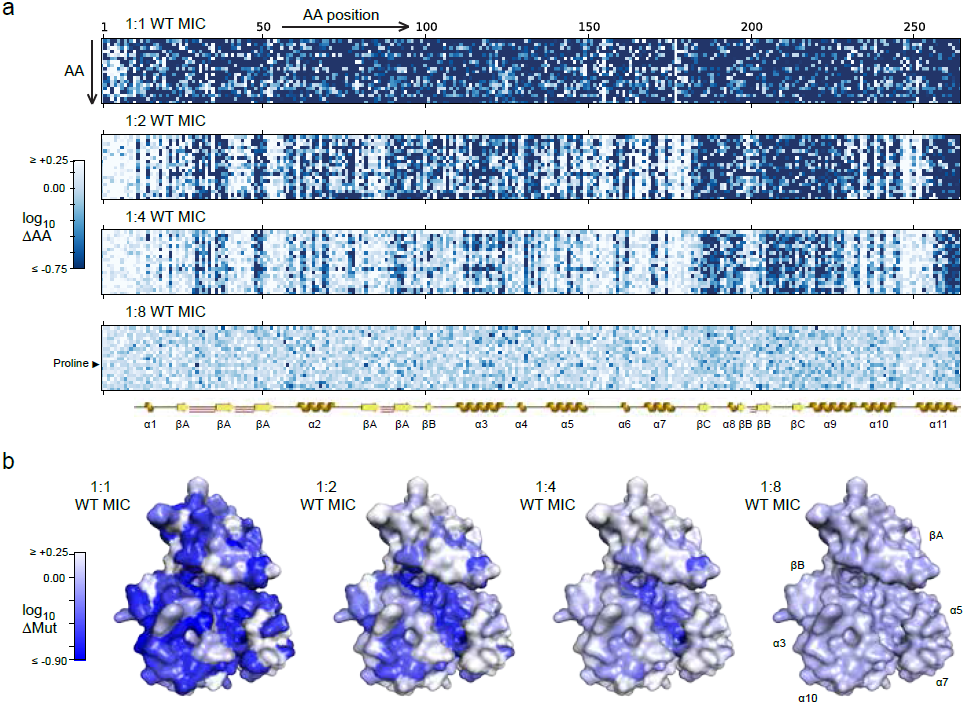
Mutational scanning of APH(3’)II using G418 selection. a. Visual representation of the abundance of each amino acid (ΔAA) at each position after selection at four different concentrations. The color in each matrix entry corresponds to the change relative to the input library. The known secondary structure of APH(3’)II is shown for reference, including alpha helices (gold), beta sheet strands (yellow) and beta hairpins (red; the first 10 residues are unstructured). Note that G418 did not provide an effective selection pressure at its 1:8 WT MIC.
b. Projections of the observed changes in the abundance of mutant amino acids (ΔMut) onto a crystal structure of APH(‘3)II in complex with kanamycin (PDB accession: 1ND4; the kanamycin substrate is not show). Numerical values are provided in Supplementary Data 2.

**Supplementary Figure 8.**
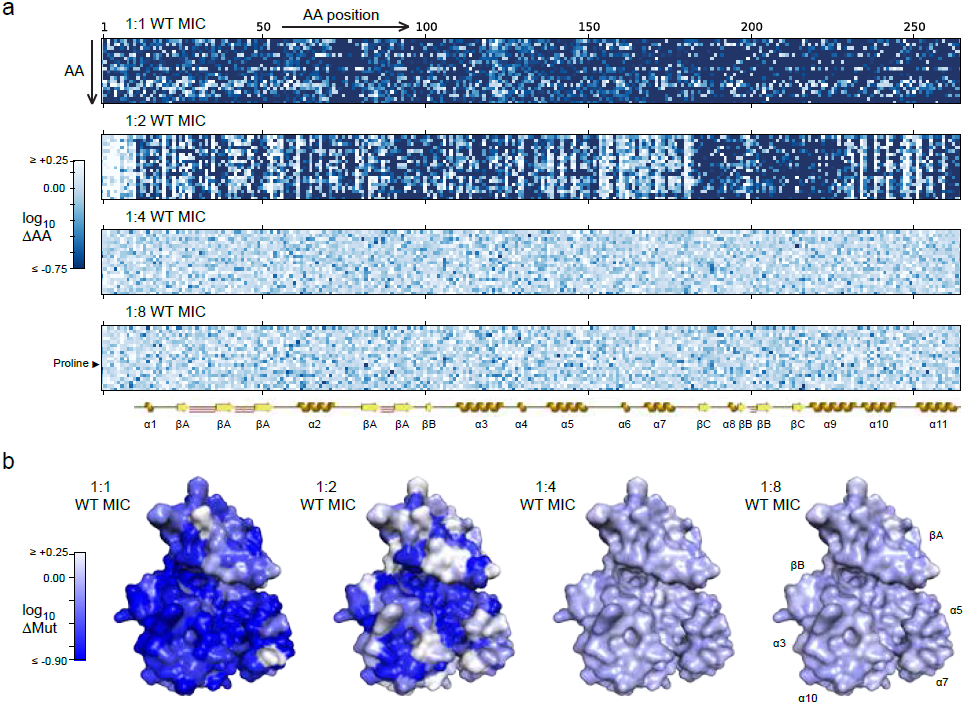
Mutational scanning of APH(3’)II using amikacin selection. a. Visual representation of the abundance of each amino acid (ΔAA) at each position after selection at four different concentrations. The color in each matrix entry corresponds to the change relative to the input library. The known secondary structure of APH(3’)II is shown for reference, including alpha helices (gold), beta sheet strands (yellow) and beta hairpins (red; the first 10 residues are unstructured). Note that amikacin did not provide an effective selection pressure at its 1:4 and 1:8 WT MICs.
b. Projections of the observed changes in the abundance of mutant amino acids (ΔMut) onto a crystal structure of APH(‘3)II in complex with kanamycin (PDB accession: 1ND4; the kanamycin substrate is not show). Numerical values are provided in Supplementary Data 2.

**Supplementary Figure 9.**
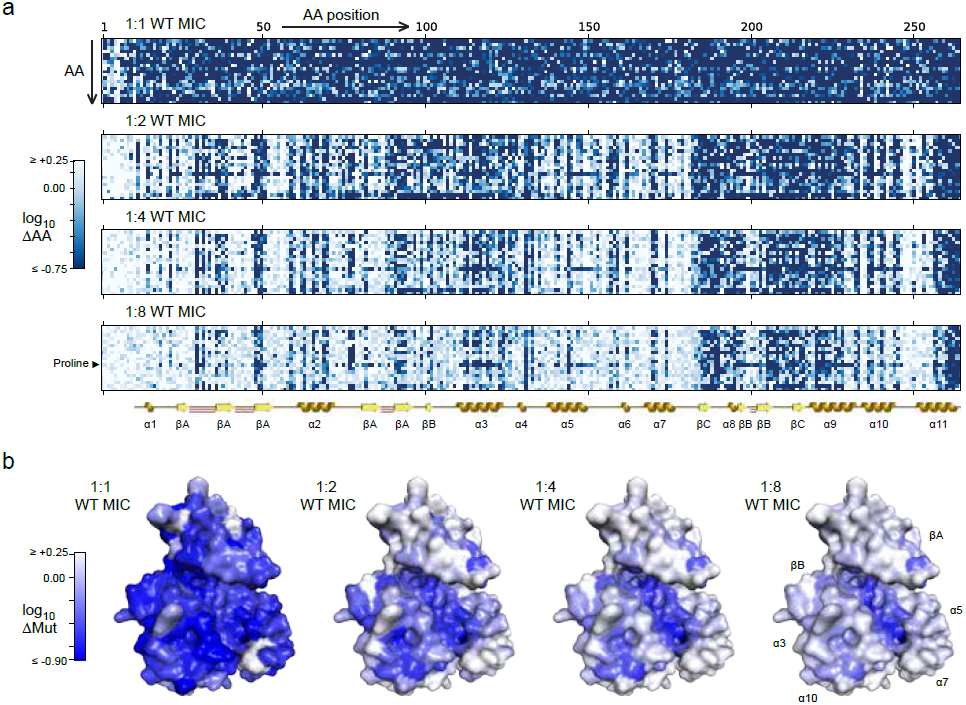
Mutational scanning of APH(3’)II using neomycin selection. a. Visual representation of the abundance of each amino acid (ΔAA) at each position after selection at four different concentrations. The color in each matrix entry corresponds to the change relative to the input library. The known secondary structure of APH(3’)II is shown for reference, including alpha helices (gold), beta sheet strands (yellow) and beta hairpins (red; the first 10 residues are unstructured).
b. Projections of the observed changes in the abundance of mutant amino acids (ΔMut) onto a crystal structure of APH(‘3)II in complex with kanamycin (PDB accession: 1ND4; the kanamycin substrate is not show). Numerical values are provided in Supplementary Data 2.

**Supplementary Figure 10.**
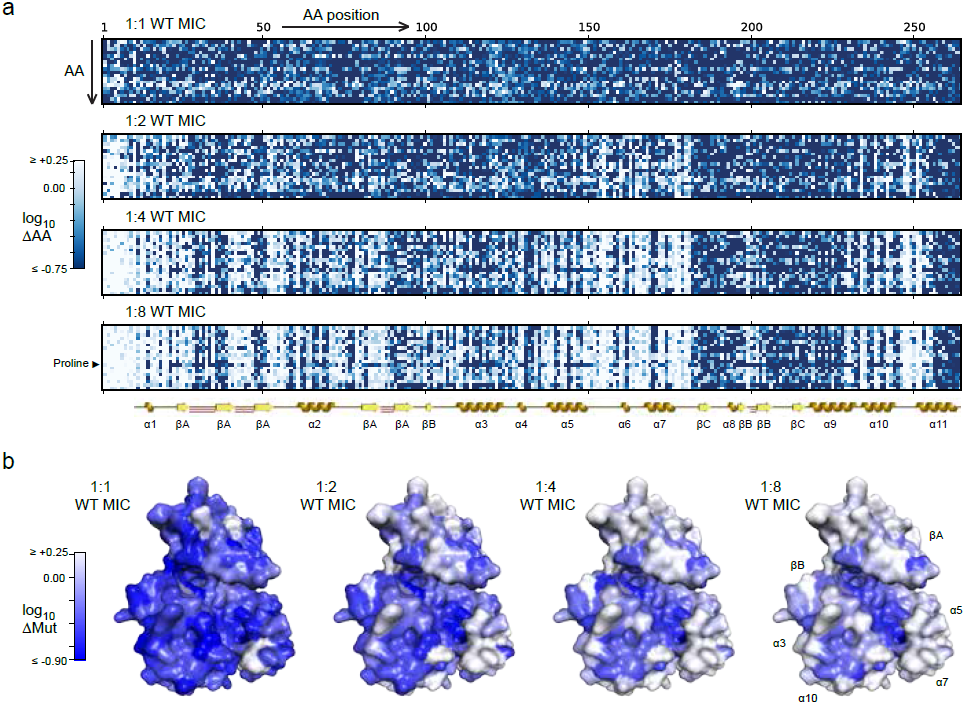
Mutational scanning of APH(3’)II using paromomycin selection. a. Visual representation of the abundance of each amino acid (ΔAA) at each position after selection at four different concentrations. The color in each matrix entry corresponds to the change relative to the input library. The known secondary structure of APH(3’)II is shown for reference, including alpha helices (gold), beta sheet strands (yellow) and beta hairpins (red; the first 10 residues are unstructured).
b. Projections of the observed changes in the abundance of mutant amino acids (ΔMut) onto a crystal structure of APH(‘3)II in complex with kanamycin (PDB accession: 1ND4; the kanamycin substrate is not show). Numerical values are provided in Supplementary Data 2.

**Supplementary Figure 11.**
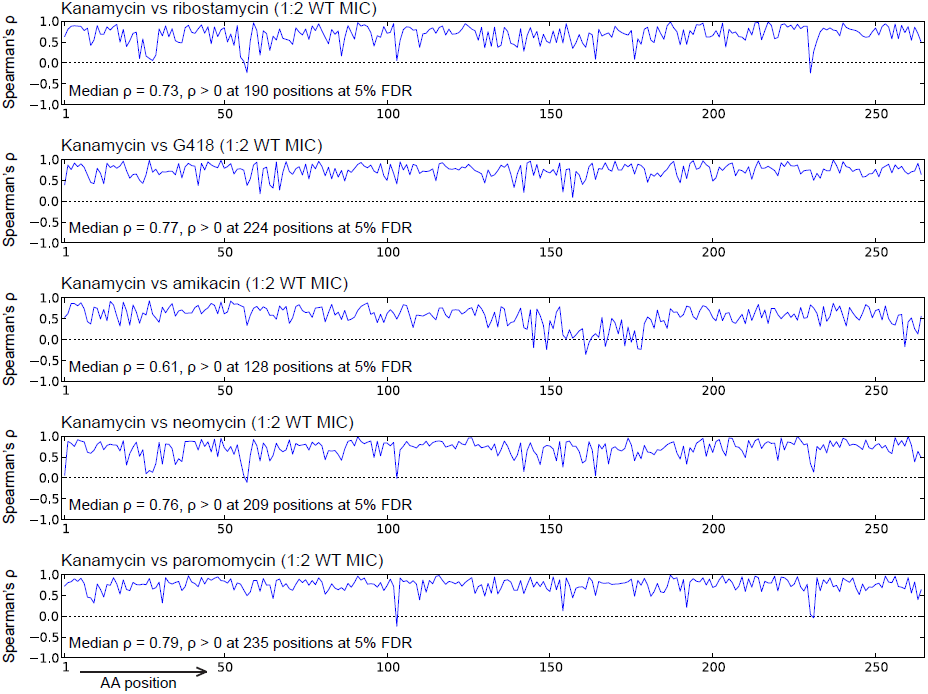
Plots of the rank correlation coefficients between the 20 possible amino acids at each position along APH(3’)II, after ranking them by the changes in their relative abundances under selection with each indicated pair of aminoglycosides at 1:2 WT MICs.

**Supplementary Figure 12.**
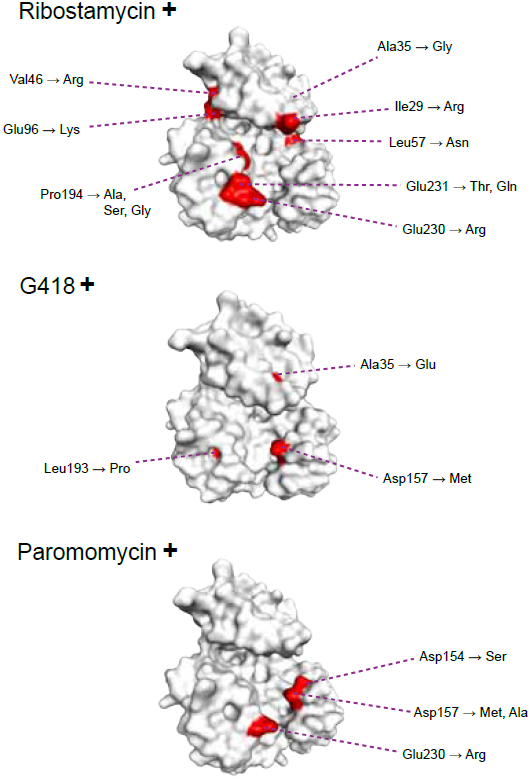
Positions and identities of amino acid substitutions that appear to be specifically tolerated or favored under selection with the indicated aminoglycoside at its 1:1 WT MIC, but not under selection with kanamycin at its 1:2 WT MIC. See also Supplementary Tables 1 and 2.

**Supplementary Table 1.** Candidate specificity-modifying substitutions.

| Selection conditions | Substitutions specifically tolerated under selection with kanamycin | Substitutions specifically tolerated under selection with second aminoglycoside |
| --- | --- | --- |
| G418 at 1:1 WT MIC, Kanamycin at 1:2 WT MIC | n/a | Glu35, Met157, Pro193 |
| G418 at 1:2 WT MIC, Kanamycin at 1:1 WT MIC | None | n/a |
| G418 at 1:2 WT MIC, Kanamycin at 1:2 WT MIC | Glu159, Ile192, Ser227, Arg231, Lys231 | Glu28, Glu31, Phe55, Cys153, Gly154, Thr154, Gln157, Met157, Arg157, Pro160, Pro193, Thr198 |
| Paromomycin at 1:1 WT MIC, Kanamycin at 1:2 WT MIC | n/a | Ser154, Met157, Ala157, Arg230 |
| Paromomycin at 1:2 WT MIC, Kanamycin at 1:1 WT MIC | Lys31 | n/a |
| Paromomycin at 1:2 WT MIC, Kanamycin at 1:2 WT MIC | Lys31, Arg31, Glu159, Ile192, Ser227, Arg231, Phe231, Lys231, Trp231 | Glu31, Ser103, Ala103, Gln154, Gln157, Leu157, Met157, Gly157, Arg230, Asn230 |
| Ribostamycin at 1:1 WT MIC, Kanamycin at 1:2 WT MIC | n/a | Arg29, Gly35, Arg46, Asn57, Lys96, Ser194, Gly194, Ala194, Arg230, Thr231, Gln231 |
| Ribostamycin at 1:2 WT MIC, Kanamycin at 1:1 WT MIC | Ser35, His55 | n/a |
| Ribostamycin at 1:2 WT MIC, Kanamycin at 1:2 WT MIC | Asn27, Ser27, Tyr28, Leu28, Trp35, Gly39, Tyr55, Gln104, Ile192, Tyr255 | Leu27, Ser29, Arg29, Arg46, Asn57, Arg96, Ser101, Arg103, Gly194, Ala194, Arg230, Asn230, Ser231, Gln231, Thr231 |
| Neomycin at 1:1 WT MIC, Kanamycin at 1:2 WT MIC | n/a | None |
| Neomycin at 1:2 WT MIC, Kanamycin at 1:1 WT MIC | His55 | n/a |
| Neomycin at 1:2 WT MIC, Kanamycin at 1:2 WT MIC | Tyr28, Arg31, Pro33, Trp35, Gln104, Glu159, Pro176, Pro178, Val179, Ile192, Arg231, Lys231, Phe231, Trp231 | Asp29, Ser29, Glu31, Trp37, Arg46, Asn56, Asn57, Lys96, Asp194, Gly194 |
| Amikacin at 1:1 WT MIC, Kanamycin at 1:2 WT MIC | n/a | None |
| Amikacin at 1:2 WT MIC, Kanamycin at 1:1 WT MIC | Ile79, Ser127, Phe137, Arg137, Tyr137, Val141, Asn149, Ile149, Phe153, Phe158, Tyr158, Trp158, Tyr163, Phe163, Glu163, Met163, Leu167, Leu173, Asn177, Trp177, Ser177, Ala177, Asp177, Gln177, Leu177, Gly177, Thr177, Tyr177, Phe177, Tyr178, Phe178, Trp256 | n/a |
| Amikacin at 1:2 WT MIC, Kanamycin at 1:2 WT MIC | Phe137, Tyr137, Arg137, Val141, Leu149, Lys149, Asn149, Ile149, Asn151, His151, Phe153, Leu155, Phe158, Tyr158, Trp158, Tyr163, Glu163, Met163, His165, Glu165, Tyr165, Trp165, Phe165, Leu167, Leu168, Glu173, Gln173, Cys175, Pro176, Asn177, Gln177, Ser177, Ala177, Asp177, Tyr177, Leu177, Gly177, Trp177, Thr177, Phe177, Tyr178, Val178, Trp178, His178, Phe178, Gln178, Val179, Thr219, Cys225, Tyr230, Lys249, Try255, Trp256, Tyr256 | Glu35, Val145, Met145, Ile145, Leu145, Cys145, Tyr148, Lys156, Gln157, Glu157, Ile157, Tyr162, Leu162, Trp162, Met162, Ile162, Phe162, Asp168, Ile170, Glu171, Ile174, Arg255, Gln259, Phe259, Thr259, Ala259, Ser259, Trp259, Asp259, His259 |

**Supplementary Table 2.**
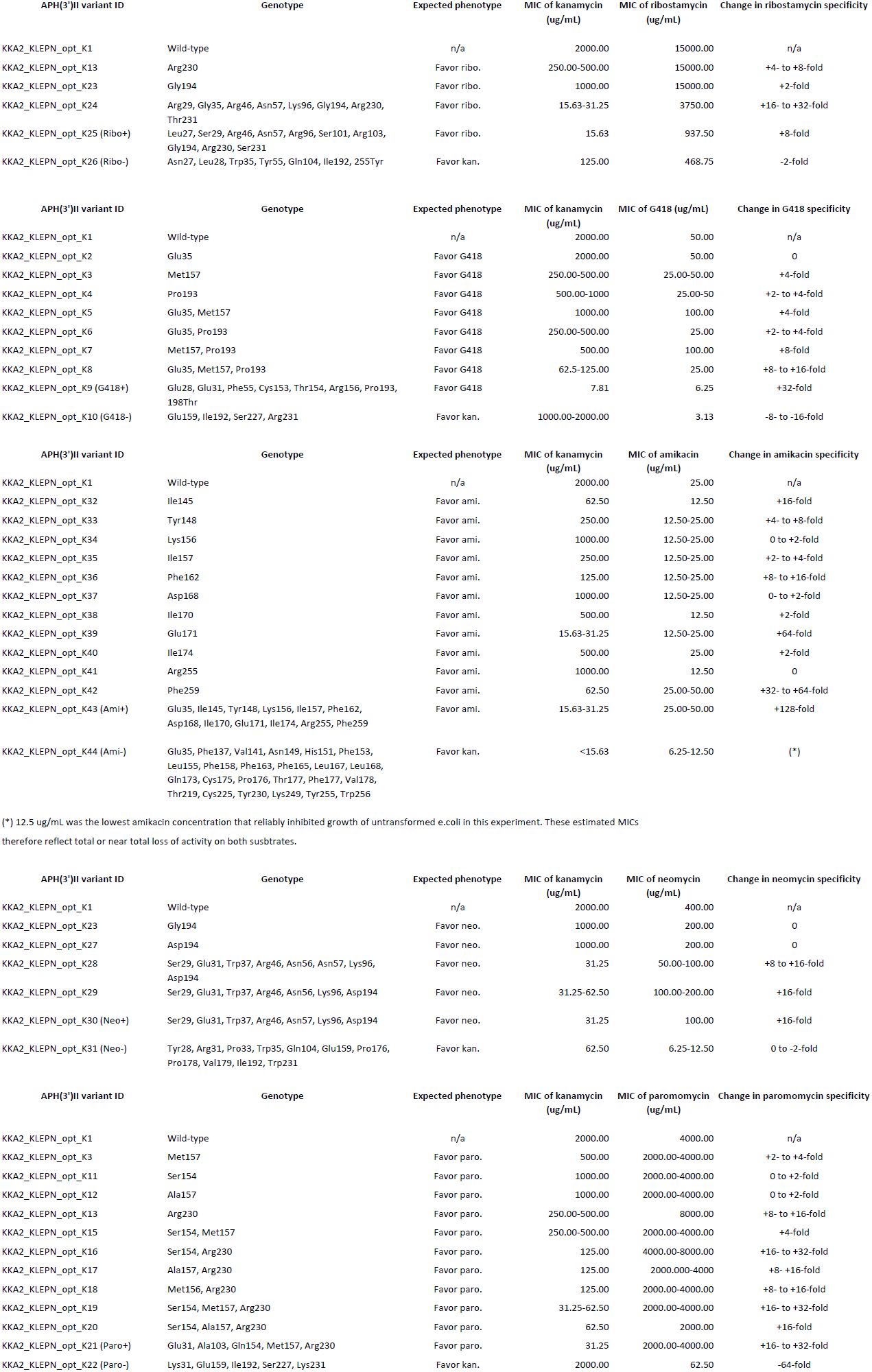
Minimum inhibitory concentrations (MICs) MICs were determined from the A_600 from 2-3 independent cultures using 2-fold dilution series. The MIC estimated from all matched cultures were identical within the resolution of the assay, expect where a range is shown. Note that these MICs are higher than those established for selection immediately following transformation (e.g. Supplementary Fig. 2) due to differences in recovery conditions.

| APH(3')II variant ID | Genotype | Expected phenotype | MIC of kanamycin (ug/mL) | MIC of ribostamycin (ug/mL) | Change in ribostamycin specificity |
| --- | --- | --- | --- | --- | --- |
| KKA2_KLEPN_opt_K1 | Wild-type | n/a | 2000.00 | 15000.00 | n/a |
| KKA2_KLEPN_opt_K13 | Arg230 | Favor ribo. | 250.00-500.00 | 15000.00 | +4- to +8-fold |
| KKA2_KLEPN_opt_K23 | Gly194 | Favor ribo. | 1000.00 | 15000.00 | +2-fold |
| KKA2_KLEPN_opt_K24 | Arg29, Gly35, Arg46, Asn57, Lys96, Gly194, Arg230, Thr231 | Favor ribo. | 15.63-31.25 | 3750.00 | +16- to +32-fold |
| KKA2_KLEPN_opt_K25 (Ribo+) | Leu27, Ser29, Arg46, Asn57, Arg96, Ser101, Arg103, Gly194, Arg230, Ser231 | Favor ribo. | 15.63 | 937.50 | +8-fold |
| KKA2_KLEPN_opt_K26 (Ribo-) | Asn27, Leu28, Trp35, Tyr55, Gln104, Ile192, 255Tyr | Favor kan. | 125.00 | 468.75 | -2-fold |

| APH(3')II variant ID | Genotype | Expected phenotype | MIC of kanamycin (ug/mL) | MIC of G418 (ug/mL) | Change in G418 specificity |
| --- | --- | --- | --- | --- | --- |
| KKA2_KLEPN_opt_K1 | Wild-type | n/a | 2000.00 | 50.00 | n/a |
| KKA2_KLEPN_opt_K2 | Glu35 | Favor G418 | 2000.00 | 50.00 | 0 |
| KKA2_KLEPN_opt_K3 | Met157 | Favor G418 | 250.00-500.00 | 25.00-50.00 | +4-fold |
| KKA2_KLEPN_opt_K4 | Pro193 | Favor G418 | 500.00-1000 | 25.00-50 | +2- to +4-fold |
| KKA2_KLEPN_opt_K5 | Glu35, Met157 | Favor G418 | 1000.00 | 100.00 | +4-fold |
| KKA2_KLEPN_opt_K6 | Glu35, Pro193 | Favor G418 | 250.00-500.00 | 25.00 | +2- to +4-fold |
| KKA2_KLEPN_opt_K7 | Met157, Pro193 | Favor G418 | 500.00 | 100.00 | +8-fold |
| KKA2_KLEPN_opt_K8 | Glu35, Met157, Pro193 | Favor G418 | 62.5-125.00 | 25.00 | +8- to +16-fold |
| KKA2_KLEPN_opt_K9 (G418+) | Glu28, Glu31, Phe55, Cys153, Thr154, Arg156, Pro193, 198Thr | Favor G418 | 7.81 | 6.25 | +32-fold |
| KKA2_KLEPN_opt_K10 (G418-) | Glu159, Ile192, Ser227, Arg231 | Favor kan. | 1000.00-2000.00 | 3.13 | -8- to -16-fold |

| APH(3')II variant ID | Genotype | Expected phenotype | MIC of kanamycin (ug/mL) | MIC of amikacin (ug/mL) | Change in amikacin specificity |
| --- | --- | --- | --- | --- | --- |
| KKA2_KLEPN_opt_K1 | Wild-type | n/a | 2000.00 | 25.00 | n/a |
| KKA2_KLEPN_opt_K32 | Ile145 | Favor ami. | 62.50 | 12.50 | +16-fold |
| KKA2_KLEPN_opt_K33 | Tyr148 | Favor ami. | 250.00 | 12.50-25.00 | +4- to +8-fold |
| KKA2_KLEPN_opt_K34 | Lys156 | Favor ami. | 1000.00 | 12.50-25.00 | 0 to +2-fold |
| KKA2_KLEPN_opt_K35 | Ile157 | Favor ami. | 250.00 | 12.50-25.00 | +2- to +4-fold |
| KKA2_KLEPN_opt_K36 | Phe162 | Favor ami. | 125.00 | 12.50-25.00 | +8- to +16-fold |
| KKA2_KLEPN_opt_K37 | Asp168 | Favor ami. | 1000.00 | 12.50-25.00 | 0- to +2-fold |
| KKA2_KLEPN_opt_K38 | Ile170 | Favor ami. | 500.00 | 12.50 | +2-fold |
| KKA2_KLEPN_opt_K39 | Glu171 | Favor ami. | 15.63-31.25 | 12.50-25.00 | +64-fold |
| KKA2_KLEPN_opt_K40 | Ile174 | Favor ami. | 500.00 | 25.00 | +2-fold |
| KKA2_KLEPN_opt_K41 | Arg255 | Favor ami. | 1000.00 | 12.50 | 0 |
| KKA2_KLEPN_opt_K42 | Phe259 | Favor ami. | 62.50 | 25.00-50.00 | +32- to +64-fold |
| KKA2_KLEPN_opt_K43 (Ami+) | Glu35, Ile145, Tyr148, Lys156, Ile157, Phe162, Asp168, Ile170, Glu171, Ile174, Arg255, Phe259 | Favor ami. | 15.63-31.25 | 25.00-50.00 | +128-fold |
| KKA2_KLEPN_opt_K44 (Ami-) | Glu35, Phe137, Val141, Asn149, His151, Phe153, Leu155, Phe158, Phe163, Phe165, Leu167, Leu168, Gln173, Cys175, Pro176, Thr177, Phe177, Val178, Thr219, Cys225, Tyr230, Lys249, Tyr255, Trp256 | Favor kan. | <15.63 | 6.25-12.50 | (*) |

| APH(3')II variant ID | Genotype | Expected phenotype | MIC of kanamycin (ug/mL) | MIC of neomycin (ug/mL) | Change in neomycin specificity |
| --- | --- | --- | --- | --- | --- |
| KKA2_KLEPN_opt_K1 | Wild-type | n/a | 2000.00 | 400.00 | n/a |
| KKA2_KLEPN_opt_K23 | Gly194 | Favor neo. | 1000.00 | 200.00 | 0 |
| KKA2_KLEPN_opt_K27 | Asp194 | Favor neo. | 1000.00 | 200.00 | 0 |
| KKA2_KLEPN_opt_K28 | Ser29, Glu31, Trp37, Arg46, Asn56, Asn57, Lys96, Asp194 | Favor neo. | 31.25 | 50.00-100.00 | +8 to +16-fold |
| KKA2_KLEPN_opt_K29 | Ser29, Glu31, Trp37, Arg46, Asn56, Lys96, Asp194 | Favor neo. | 31.25-62.50 | 100.00-200.00 | +16-fold |
| KKA2_KLEPN_opt_K30 (Neo+) | Ser29, Glu31, Trp37, Arg46, Asn57, Lys96, Asp194 | Favor neo. | 31.25 | 100.00 | +16-fold |
| KKA2_KLEPN_opt_K31 (Neo-) | Tyr28, Arg31, Pro33, Trp35, Gln104, Glu159, Pro176, Pro178, Val179, Ile192, Trp231 | Favor kan. | 62.50 | 6.25-12.50 | 0 to -2-fold |

| APH(3')II variant ID | Genotype | Expected phenotype | MIC of kanamycin (ug/mL) | MIC of paromomycin (ug/mL) | Change in paromomycin specificity |
| --- | --- | --- | --- | --- | --- |
| KKA2_KLEPN_opt_K1 | Wild-type | n/a | 2000.00 | 4000.00 | n/a |
| KKA2_KLEPN_opt_K3 | Met157 | Favor paro. | 500.00 | 2000.00-4000.00 | +2- to +4-fold |
| KKA2_KLEPN_opt_K11 | Ser154 | Favor paro. | 1000.00 | 2000.00-4000.00 | 0 to +2-fold |
| KKA2_KLEPN_opt_K12 | Ala157 | Favor paro. | 1000.00 | 2000.00-4000.00 | 0 to +2-fold |
| KKA2_KLEPN_opt_K13 | Arg230 | Favor paro. | 250.00-500.00 | 8000.00 | +8- to +16-fold |
| KKA2_KLEPN_opt_K15 | Ser154, Met157 | Favor paro. | 250.00-500.00 | 2000.00-4000.00 | +4-fold |
| KKA2_KLEPN_opt_K16 | Ser154, Arg230 | Favor paro. | 125.00 | 4000.00-8000.00 | +16- to +32-fold |
| KKA2_KLEPN_opt_K17 | Ala157, Arg230 | Favor paro. | 125.00 | 2000.000-4000 | +8- +16-fold |
| KKA2_KLEPN_opt_K18 | Met156, Arg230 | Favor paro. | 125.00 | 2000.00-4000.00 | +8- to +16-fold |
| KKA2_KLEPN_opt_K19 | Ser154, Met157, Arg230 | Favor paro. | 31.25-62.50 | 2000.00-4000.00 | +16- to +32-fold |
| KKA2_KLEPN_opt_K20 | Ser154, Ala157, Arg230 | Favor paro. | 62.50 | 2000.00 | +16-fold |
| KKA2_KLEPN_opt_K21 (Paro+) | Glu31, Ala103, Gln154, Met157, Arg230 | Favor paro. | 31.25 | 2000.00-4000.00 | +16- to +32-fold |
| KKA2_KLEPN_opt_K22 (Paro-) | Lys31, Glu159, Ile192, Ser227, Lys231 | Favor kan. | 2000.00 | 62.50 | -64-fold |

